# *BinderSpace*: A Package for Sequence Space Analyses for Datasets of Affinity-Selected Oligonucleotides and Peptide-Based Molecules

**DOI:** 10.1101/2023.02.15.528758

**Authors:** Payam Kelich, Huanhuan Zhao, Lela Vuković

**Author notes:** P. K. and H. Z. contributed equally.

## Abstract

Discovery of target-binding molecules, such as aptamers and peptides, is usually performed with the use of high-throughput experimental screening methods. These methods typically generate large datasets of sequences of target-binding molecules, which can be enriched with high affinity binders. However, the identification of the highest affinity binders from these large datasets often requires additional low-throughput experiments or other approaches. Bioinformatics-based analyses could be helpful to better understand these large datasets and identify the parts of the sequence space enriched with high affinity binders. BinderSpace is an open-source Python package that performs motif analysis, sequence space visualization, clustering analyses, and sequence extraction from clusters of interest. The motif analysis, resulting in text-based and visual output of motifs, can also provide heat maps of previously measured user-defined functional properties for all the motif-containing molecules. Users can also run principal component analysis (PCA) and t-distributed stochastic neighbor embedding (t-SNE) analyses on whole datasets and on motif-related subsets of the data. Functionally important sequences can also be highlighted in the resulting PCA and t-SNE maps. If points (sequences) in two-dimensional maps in PCA or t-SNE space form clusters, users can perform clustering analyses on their data, and extract sequences from clusters of interest. We demonstrate the use of BinderSpace on a dataset of oligonucleotides binding to single-wall carbon nanotubes in the presence and absence of a bioanalyte, and on a dataset of cyclic peptidomimetics binding to bovine carbonic anhydrase protein.

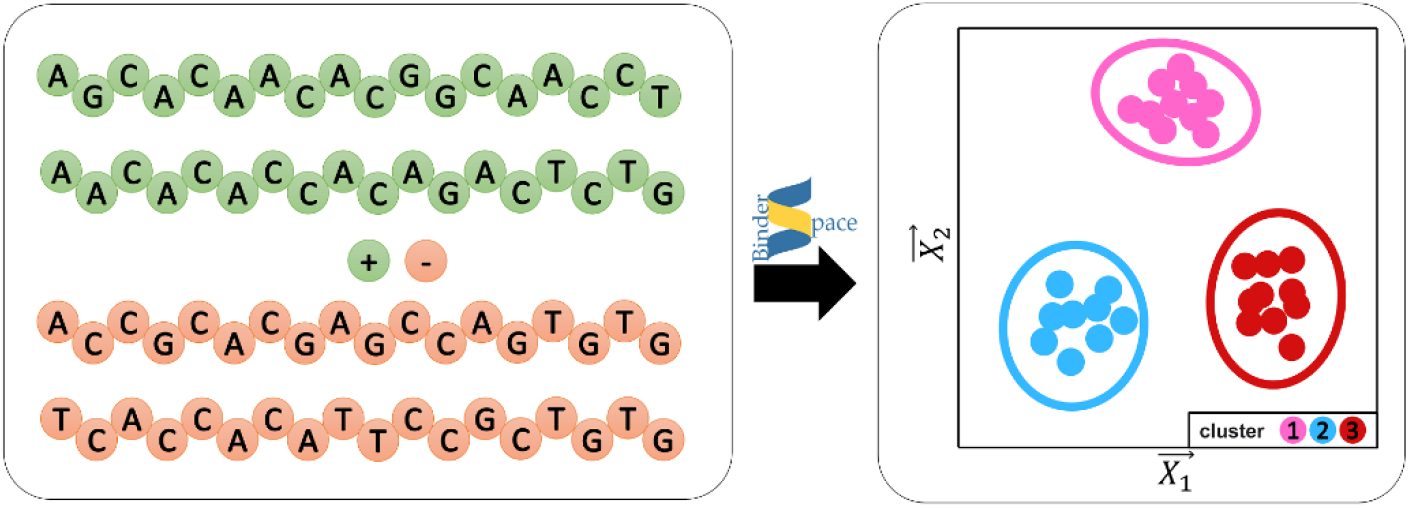

## Introduction

One of the essential steps in the development of new therapeutics and research tools is the discovery of molecules that bind with high affinity to target molecules^1,2^. The discovery of such molecules is challenging, especially if they are also required to simultaneously have selective binding to the target, and potentially also lead to another functionally important activity, such as emitting optical signals or inducing a targeted protein degradation^3^. A widely used approach for discovery of target-binding molecules relies on synthesis and screening of combinatorial libraries of molecules, which can be standalone or displayed on other entities, such as oligonucleotides or phages. In display experiments, the oligonucleotide sequence or the phage genome sequence can be read out by the next generation sequencing and provide unique encoding to each molecule in the library. The encoded libraries of molecules can be simultaneously screened in one pot for binding to the target of interest, and this process has been shown to result in a selection of high affinity binders from the original library.

The types of molecules comprising the libraries can include the oligonucleotides themselves, functionalized nucleic acid polymers^4^, small molecules^5,6^, linear, cyclic or chemically modified peptides^7–11^, and novel hybrid molecules^12^. A method for selecting target binders from libraries of oligonucleotides of varying sequences^13,14^ is called SELEX (systematic evolution of ligands by exponential enrichment). This method can be performed in multiple rounds, where in each round, high affinity binders are getting more and more enriched in the set of oligonucleotides present in the solution and binding to the target. SELEX experiments typically result in a dataset of oligonucleotide sequences enriched in selection for target binding. The targets in SELEX experiments can include small molecules^15^, cancer cells^16^, carbon nanotubes^17^. Once oligonucleotides with high affinity binding to the target, also called aptamers, are identified, they can be used for various applications.

Selection-based high-throughput screening of libraries of peptide-based molecules has also been developed over the last few decades. As mentioned above, widely used technologies for peptide-based ligand discovery are using experimental display technologies^18–20^ (e.g. phage, messenger RNA display). In these techniques, vast libraries of peptide-based molecules (as high as 10^10^) can be synthesized and assayed for binding to target proteins. Each ligand library is typically based on a single structural scaffold that contains peptide segments with fixed and variable amino acid positions, sometimes chemically modified by synthetic fragments^7–9,21^. Similar to SELEX, selection of peptides from phage or mRNA display libraries typically results in datasets of *thousands* of sequences of target-binding molecules. The obtained datasets are likely enriched with molecules with high affinity to targets. However, the definitive discovery of high affinity ligands requires additional low throughput experiments measuring the equilibrium dissociation constant *K_D_* for each selected molecule (oligonucleotide or peptide/peptidomimetic) and its target. These experiments are typically low-throughput and can be performed for a limited number of selected molecules. Thus, the identification of molecules with the highest affinity to targets requires further experiments or other approaches.

With the advent of artificial intelligence (AI) approaches, there is a large interest in using experimental selection datasets for training machine learning (ML) models to predict oligonucleotides, peptide-based and other molecules with high affinity for targets^4,22^ or the highest ability to impart functional response to the target, such as the induction of a fluorescence signal^23–25^. Bioinformatics analyses can be used to understand sequence composition of experimental datasets and also to assess the sequence patterns in sets of molecules predicted to have the highest affinity for targets by the ML models. For example, sequence motif analyses can determine the motifs that are enriched in experimental datasets, which can be useful because sequence motifs are known to be important in sequence affinity for the target^26^. Furthermore, comparative sequence motif analyses of control and selection datasets for one or multiple targets could be a helpful guide for choosing candidate molecules for follow up experiments that test in finer detail the candidates’ target binding or functional activities. Bioinformatics analyses for motif discovery in DNA, protein, and peptide datasets have been developed and successfully applied to identify short DNA sequence segments (10-30 bp) implicated in gene regulation, for developing new aptamers^22^, for discovery of antimicrobial or anticancer peptides^27^, and many other applications. Widely used codes for motif discovery include MEME^28^ suite webserver and MERCI^29^ tool written in Perl. These codes have been optimized to perform the motif discovery tasks for longer sequences and may not have the options needed to analyze selection datasets, such as the identification of very short motifs in selection libraries of smaller molecules and the simultaneous analysis of control and selection datasets.

Another useful bioinformatics analysis of sequence composition in selection datasets is the visual analysis in spaces of reduced dimensionality. Commonly used analyses include the principal component analyses (PCA) and t-distributed stochastic neighbor embedding (t-SNE), which can visualize the local and global structures within data. These analyses have been applied to large scale sequence datasets in biology, including the DNA methylation data^30^ and single-cell transcriptomes^31^. These analyses can be accompanied by the clustering approaches applied to data visualized in the reduced dimensionality spaces. The clustering approaches could be useful to identify which sequences are related to each other in these reduced dimensionality spaces. The sequences found to belong to the same clusters in the reduced dimensionality spaces as the highly functional sequences may also share their functional properties and could thus be of interest in the search for molecules with the highest affinity for targets and the highest ability to impart functional response to the target.

To the best of our knowledge, there is currently no single toolkit package that integrates the motif discovery and reduced dimensionality sequence space visualizations for analyses of oligonucleotide and peptide-based selection datasets. Here, we developed a python package called *BinderSpace*, which is designed to examine and understand the sequence composition of datasets of oligonucleotides and peptide/peptidomimetic molecules obtained by selection processes in a fast and efficient manner. *BinderSpace* package can perform motif analyses of DNA, RNA, or peptide sequences in datasets, visualize sequences that contain specific motifs in reduced dimensionality spaces, and perform cluster analyses for whole sets or subsets of sequences. We demonstrate the application of our package to two types of datasets, namely, the datasets of oligonucleotides selected for binding to single-wall carbon nanotubes in the presence of serotonin analyte^17,24^ and datasets of cyclic peptidomimetics selected for binding to bovine carbonic anhydrase (BCA) protein target^8^.

## Methods

Here, we describe the BinderSpace package, which analyzes datasets of related DNA and peptide-based sequences of molecules obtained in selection experiments. The analyses currently implemented in BinderSpace, summarized in **Figure 1**, include motif analysis, sequence space visualization, cluster analysis and sequence extraction from clusters of interest.

**Figure 1.**
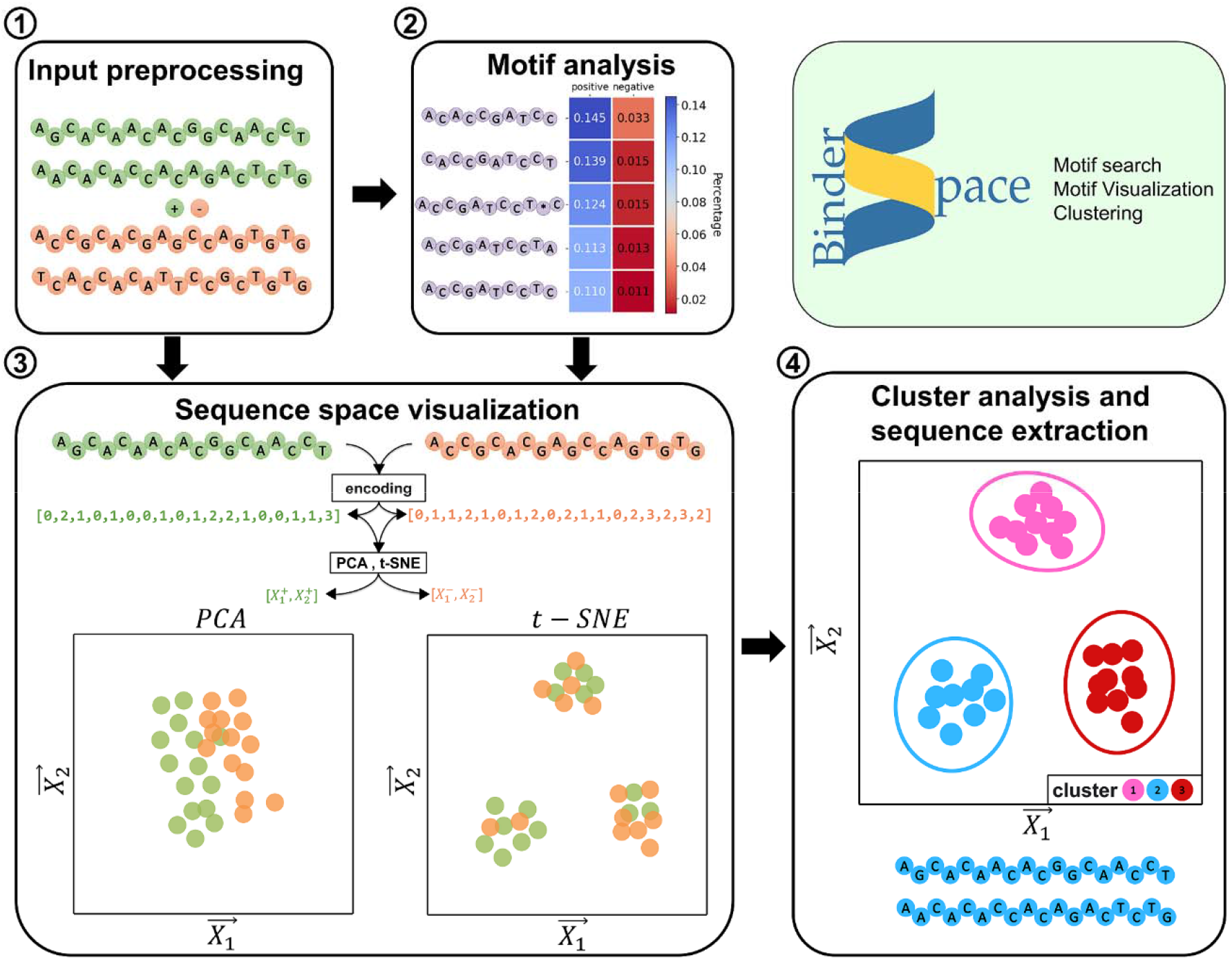
Overview of functionalities of the BinderSpace package. The inputs of the package (box 1) are DNA or peptide molecule sequences, including molecules with high-affinity binding to target (positive dataset) and molecules from control dataset assumed to have no high-affinity binding to target (negative dataset). The first output (box 2) is the analysis of motifs in positive and negative datasets. The second output (box 3) is the visual analysis of the sequence space from positive and negative datasets. The third output (box 4) is the cluster analysis of the sequenc space plots. Sequences of molecules can be extracted from clusters of interest.

### Implementation

BinderSpace is an open-source package implemented in Python3. The source code is available on GitHub and can be installed or downloaded from PyPI (Python Package Index) repository. The repository contains a folder with the code and a folder with an example analysis in the tutorial form.

### Motif Search Task for Datasets of Affinity-Selected Molecules

Motif search is the first BinderSpace task in the analysis of datasets of affinity-selected DNA or amino acid-based molecules. The datasets can contain sequences with high affinity binding to the target, which are also called positive sequences. Alternatively, the datasets can contain both positive and negative sequences, where the negative (control) sequences are assumed to have no high-affinity binding to target.

The motif search task, based on the Apriori alghorithm^32^, identifies top K motifs that are most frequent in the set of positive sequences. The output is the csv file with the list of motifs and their percentage in the set of positive sequences. When sets of both positive and negative sequences are analyzed, then the motif search returns top K motifs that are most frequent in a positive set of sequences, and low frequency in a negative set. The output of the motif search (a csv file) provides a list of motifs, ranked according to the difference between the motif’s percentage in the set of positive sequences and the motif’s percentage in the set of negative sequences. Since our code is intended to analyze genetically encoded affinity-selected molecules of the same size but different sequences, it assumes that sequence lengths for all the molecules in the dataset are the same.

The motif search task is performed by running motif_search.py from the command line, a parallelized Python 3 code. The required and optional flags for running this code from the command line are listed and described in **Table 1**. The only required flag is (-i), which defines the file name of the input dataset of positive sequences. There is an optional flag (-n) to call a second input dataset of negative (control) sequences. In practical applications, the control dataset can contain sequences with low affinity for targets of interest or sequences found to have non-specific affinity for targets of interest. Alternatively, a separate function in module.py (random_sequence function) can also provide a negative dataset by generating a defined number of random sequences that are not present in the positive dataset.

**Table 1.** Summary of options in *motif_search.py* code.

| flag | description | default | format |
| --- | --- | --- | --- |
| -i | calls the input file containing positive sequences (csv) | - | file name, required |
| -n | calls the input file containing negative sequences (csv) | - | file name |
| -o | defines the name of the output files listing the found motifs to be different from default (motifs.csv) | motifs.csv | file name |
| -c | defines if used for protein or DNA sequences | amino_acids | dna or amino_acids |
| -f | the minimal occurrence frequency for the positive sequences | 0.001 | 0<number < 1 |
| -m | the minimal motif length | 3 | integer > 1 |
| -l | the maximal motif length | the length of the longest positive sequence | integer > 1 |
| -g | maximal number of gaps in motifs | 0 | integer ≥ 0 |
| -a | maximal gap length | 1 | integer ≥ 1 |
| -p | number of processors used for running motif_search.py | 20 | integer |

The motif search task has several other options, described in **Table 1**. There is a basic flag (-c) for choosing the type of molecules in the affinity selected dataset, either DNA or peptide. The search process is defined by the flag (-f), the minimal occurrence frequency of any given motif in the positive dataset, a parameter which affects the speed of the motif search. The minimal motif length (-m) and the maximal length of the motifs (-l) can be defined, as well as the number of gaps the motifs can have (-g), and the maximal gap length (-a). Finally, our motif search code can be performed on multiple processors, and the user can define the number of processors used with (-p) flag.

### PCA and t-SNE Analyses

The second *BinderSpace* task is performing the PCA and t-SNE analyses on datasets of DNA or peptide-based molecules. These analyses can be performed on whole datasets and on motif-related subsets of the data. The code binder_space.py performs these analyses, and we recommend the user to use this code as a Jupyter notebook, in order to be able to interact with the code and make modifications as needed.

The simplest analyses in PCA and t-SNE space are analyses in two- and three-dimensional space, which allow the user to plot and visually examine the dataset in these reduced dimension spaces. The PCA and t-SNE analyses in *binder_space.py* are performed using the modules implemented in scikit-learn with parameters found to work best on the DNA dataset that we used for testing. However, the section for t-SNE analysis includes detailed guidance on hyperparameter testing that the user should perform as a routine part of the analysis. For both PCA and t-SNE analyses, the sequences of molecules in the datasets are represented either as (1 x N) matrices for peptide-based molecules containing N amino acids or (1 x M) matrices for DNA molecules containing M nucleotides (**Table S1**).

The user can decide what dataset is to be analyzed. Namely, the user can analyze whole datasets or subsets of datasets, such as subsets of sequences that contain desired motifs. The code guides the user how to perform each type of analysis and recommends how to perform optimization of the hyperparameters.

The code can also output PCA and t-SNE coordinates of the analyzed sequences for all the dimensions present in the datasets, when performing the t-SNE analysis with the regular scikit-learn library. These coordinates are written out as csv output. For large datasets, such as the DNA dataset we used for testing (>750,000 sequences), t-SNE analysis of the whole sequence space may be slow. In that case, we provide a suggestion for using cuML library for t-SNE analysis in our code, which enables only a two-dimensional analysis and currently supports a Linux-like system^33^.

### Clustering Analysis of Datasets in PCA and t-SNE Space

The next option provided by our codes, distributed as Jupyter notebooks, is to perform cluster analysis on the two-dimensional maps obtained in PCA and t-SNE analyses. Based on the shape of the data in the maps, the user can choose to perform clustering using any of four clustering methods implemented in scikit-learn library, including density-based spatial clustering of applications with noise (DBSCAN), balanced iterative reducing (Birch), Gaussian Mixture and k-means. For the DBSCAN method^34^, we advise users to test two parameters for improving the clustering results, ε, the maximum distance between two points for these points to be categorized as a cluster, and min_samples, the minimum number of samples that form a cluster. For the Birch method, the user can examine the clustering results for different values of the number of clusters to be obtained and the threshold parameter. In the Gaussian Mixture method, the clustering results may be improved by varying several parameters, including the number of clusters (n_components), covariance type, the number of iterations performed, and the method used for initial definitions of clusters (init_params). In the K-means methods, the user may change clustering results by adjusting the hyperparameter defining the number of clusters to be obtained.

### Example Datasets

Two datasets, summarized in **Table 2**, were used to demonstrate the BinderSpace options. SDNA dataset, obtained by Jeong, Landry and co-workers and reported in Ref.^17^, contains 18-nt long DNA sequences, and is divided into sub-datasets of positive and negative sequences SDNA-pos and SDNA-neg. SDNA-pos contains sequences selected for binding to single walled carbon nanotubes (SWNTs) in the presence of a serotonin neurotransmitter analyte, and SDNA-neg contains sequences selected for binding to SWNTs only (control sequences). The dataset of affinity-selected peptidomimetics BCA-pos and BCA-neg, initially obtained in experiments by Ekanayake, Derda and co-workers and described in Ref.^8^, contains 1,3-diketone bearing macrocyclic peptides (DKMP), formed of peptide fragments that vary in sequence covalently bound to 1,3-diketone group fragments. The molecules in this dataset were selected for binding to bovine carbonic anhydrase (BCA), identified by panning against BCA and found to be significantly enriched in panning to BCA when compared to panning against a control protein bovine serum albumin (BSA).

**Table 2.** Summary of datasets used to demonstrate *BinderSpace* use. X labels the variable positions in molecules.

| Molecule type | Dataset | Length | Sequence template | Number of sequences |
| --- | --- | --- | --- | --- |
| DNA | SDNA-pos | 18-nt | C<sub>6</sub>-X<sub>18</sub>-C<sub>6</sub> | 570,926 |
|  | SDNA-neg | 18-nt | C<sub>6</sub>-X<sub>18</sub>-C<sub>6</sub> | 219,382 |
| peptidomimetic | BCA-pos | 6-aa | SXCXXXC-DKMP | 7,815 |
|  | BCA-neg | 6-aa | SXCXXXC-DKMP | 7,644 |

## Results and Examples

### Motif Search

After importing the positive and negative datasets, BinderSpace analyzes the sequences for frequently occurring motifs. Then, it outputs the observed sequence motifs and their frequencies within the positive and negative dataset. The motifs can be fully sequence-based or can contain gaps. The output includes csv files that each contain sequence motifs of a given length.

We first demonstrate the motif search functionality on SDNA and BCA datasets. The example outputs of top 5 motifs of two different lengths with and without gaps in sequences are shown in **Figure 2a-b**. The *motif_search.py* program generates a csv file output for each motif length. Motifs of all lengths larger than the user-defined minimal length for the used datasets are found within a single search. In addition to the comprehensive list of motifs and their percentages in the set of positive or negative sequences being outputted in csv format, our code can also provide a visual output of top motifs along with their associated percentages colored according to magnitudes (**Figure 2a-b**), for a convenient visual evaluation on how represented the motif is in the studied dataset. **Figure 2a** also shows the capability of our code to visualize values of a functional property of interest for sequences containing each motif. For SDNA dataset, the functional property is the optical fluorescence emission change, called ΔF/F, after the addition of serotonin analyte to an aqueous sample containing single walled carbon nanotubes wrapped by DNA sequences with the listed motifs. This option can be useful to identify visually which motifs are associated with high values of the functional property of interest. The BCA dataset of peptidomimetic molecules contains lists of only four variable amino acids within molecules. Our motif search can identify 4 amino acid motifs with two gaps, as well as the 2 and 3 amino acid motifs, as shown in **Figure 2b** and **Table S4**.

**Figure 2.**
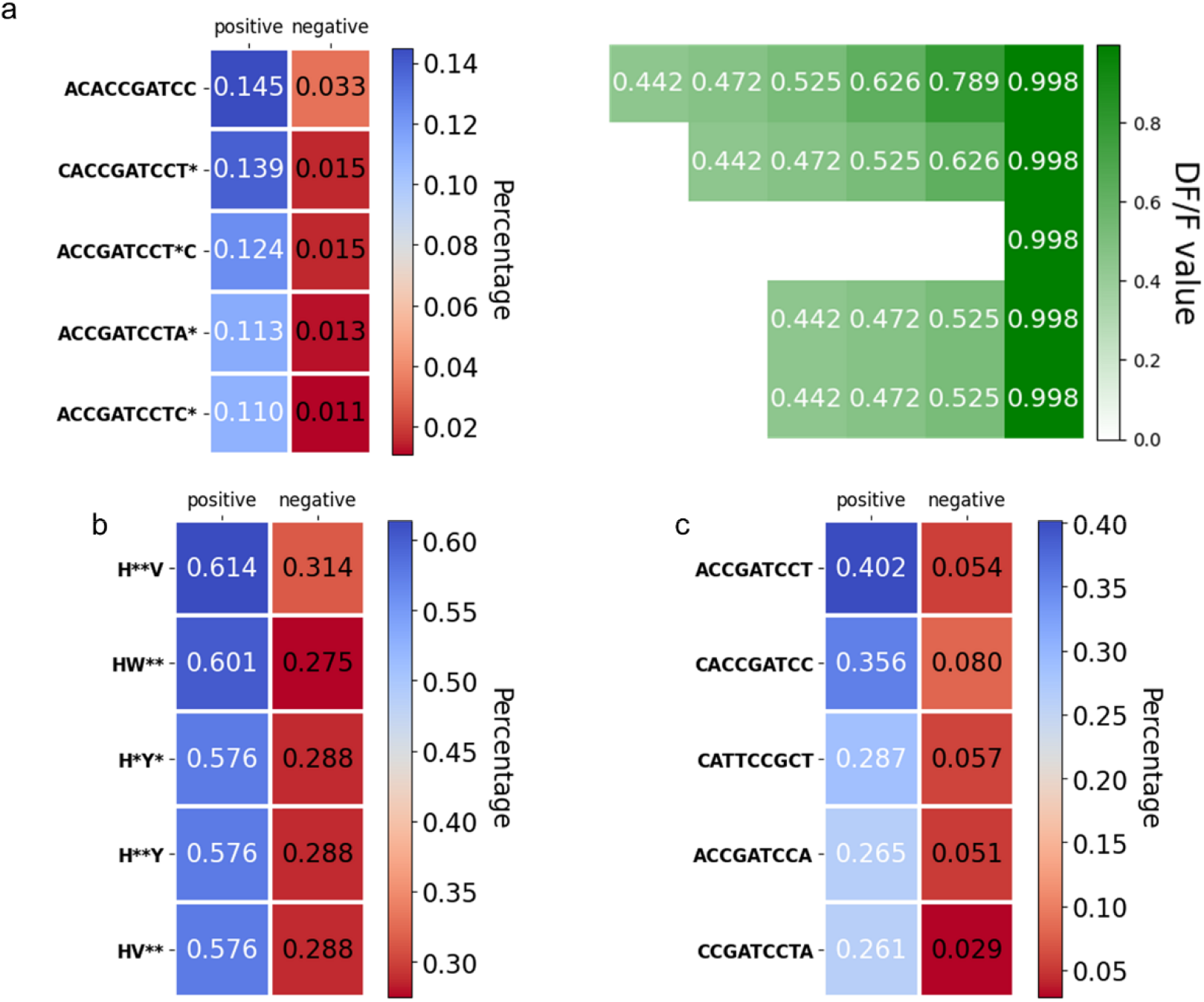
Examples of the visual output of motifs found using BinderSpace for two example datasets. The output of the motif analysis is a csv file (complete list) and a heatmap (top motifs list) in which the motifs are sorted depending on the occurrence percentage in the positive class. a) Heatmap of top 10-nt long motifs with one or no gaps from the SDNA dataset, colored according to percentages in positive and negative subsets (left). Heatmap of ΔF/F values for DNA sequences containing those motifs, obtained in experiments^24^ (right). b) Heatmap of top 4-amino acid long motifs with two gaps from BCA dataset. c) Heatmap of top 9-nt long motifs without gaps from SDNA dataset.

The running time of BinderSpace motif_search.py to find the motifs from SDNA dataset, with 790,308 sequences in total, depends on the frequency option setting (-f) and the number of gaps (-g). The running time is faster for larger frequencies and fewer gaps (**Table S2**). For example, for SDNA dataset with default frequency of 0.001 and 1 gap, the running time to generate all motifs 8 nt and longer in length was 1,128 s (18.8 min). However, when we change the frequency to 0.01%, the running time reduces to 85 s (1.42 min). The running time of BinderSpace to find motifs for BCA dataset, containing in total 15,459 amino acid sequences four amino acids in length, was always less than 3 seconds (**Table S2**).

### Examining the Sequence Space Covered by Datasets of Affinity-Selected Molecules

To visualize the sequence space of affinity-selected molecules in SDNA dataset, PCA and t-SNE analyses were performed for 18-nt DNAs initially represented as 1 x 18 encoded arrays (**Table S1**). **Figure 3a-b** shows the location of top 500 SDNA-pos and SDNA-neg dataset sequences, shown overlapped with all the SDNA-pos and SDNA-neg sequences containing C*CATTCCGCT motif, identified in a functionally interesting sequence in a previous study^17^ (ACGCCAACACATTCCGCT, labeled as E6#6, the sixth most abundant sequence in the sixth round of selection for binding to serotonin analyte in the presence of the SWNT sample). The particular sequence and motif were selected because a sample of single-walled carbon nanotubes wrapped by this sequence had 169% increase in optical emission in the presence of serotonin analyte^17^, as measured for (8,6) SWNTs at the wavelength of 1195 nm. The PCA and t-SNE analyses were performed in the spaces defined by all 790,308 sequences from the SDNA dataset, but only selected sequences are visualized in those spaces for clarity. **Figure 3a-b** shows that four subsets of DNA sequences cover similar regions of the sequence space in both PCA and t-SNE analyses. Due to the high overlap, it is difficult to say which regions of PCA and t-SNE spaces are associated with sequences that bind to serotonin analyte versus sequences that do not bind to serotonin analyte.

**Figure 3.**
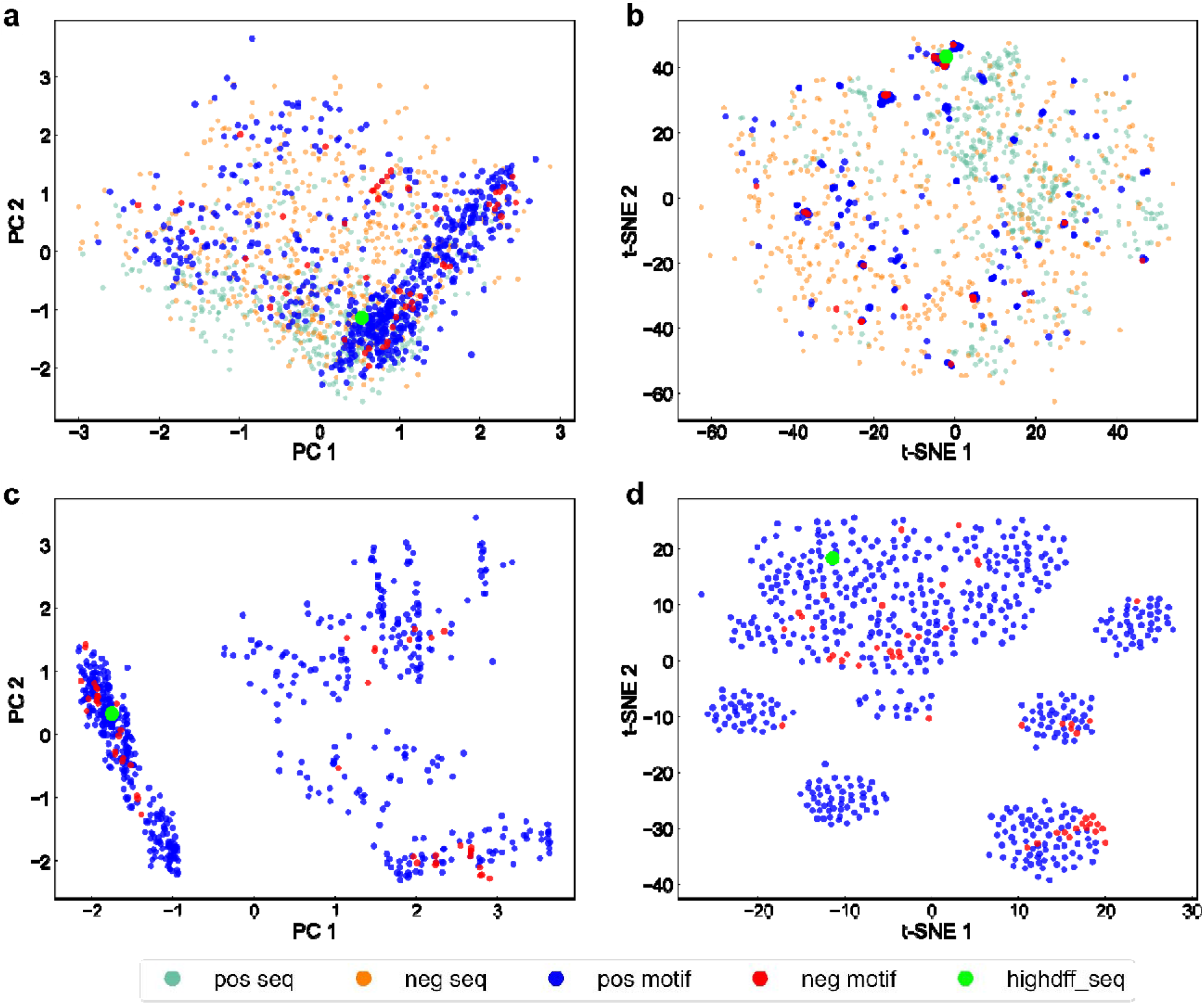
Examination of the sequence space covered by the SDNA dataset. The maps were generated by applying the PCA or t-SNE methods for dimensionality reduction to 18-nt long DNA sequences initially represented as 1 × 18 arrays. a) Comparison of top 500 sequences from SDNA-pos (orange) and SDNA-neg datasets (green), shown together with sequences containing C*CATTCCGCT motif from SDNA-pos (blue) and SDNA-neg (red) datasets. The motif was identified in ACGCCAACACATTCCGCT sequence (fluorescent green point), a functionally interesting sequence^24^. The principal component (PC) space is obtained from the complete SDNA dataset. b) Comparison of the same sequences as in panel a with t-SNE analysis. The t-SNE space is obtained from the complete SDNA dataset. c) Comparison of sequences containing C*CATTCCGCT motif from SDNA-pos (blue) an SDNA-neg (red) datasets. The PC space is obtained from SDNA sequences containing the C*CATTCCGCT motif. d) Comparison of the same sequences as in panel c with t-SNE analysis. The t-SNE space is obtained from SDNA sequences containing the C*CATTCCGCT motif.

**Figure 3c-d** shows the location of SDNA-pos and SDNA-neg sequences containing C*CATTCCGCT motif in PCA and t-SNE spaces based on the subset of SDNA sequences containing this motif. The plots have separated clusters of sequences, especially in the t-SNE space, and the clusters have different compositions of SDNA-pos and SDNA-neg sequences with the C*CATTCCGCT motif. One possible use of datasets of affinity-selected molecules, such as the SDNA dataset, is to train machine learning models to predict new high-affinity binding sequences. The analyses in **Figure 3** show the degree of separation between SDNA-pos and SDNA-neg sequences. These analyses can help us decide if ML models could be predictive and accurate. For example, since there is better separation between SDNA-pos and SDNA-neg sequences in **Figure 3c-d**, the dataset used in this figure will likely lead to better quality ML models than the datasets analyzed in **Figure 3a-b**, for which the points representing SDNA-pos and SDNA-neg sequences have poorer separation and greater overlap. However, because ML models would be based on smaller subsets of data containing specific overrepresented pattern / motif, the ML models would not generalize to predictions for sequences without this motif. Depending on the goals of the projects, the user is recommended to examine the maps generated from both complete datasets and extracted subsets to determine if predictive ML models can be obtained based on these sets of sequences.

### Clustering Analysis and Extraction of Sequences from Specific Clusters

Our results in **Figure 3c-d** show that points in the reduced dimensionality sequence space sometimes form clusters. These clusters, visible by eye, have varying compositions of sequences originating from positive and negative parts of datasets. In fact, some clusters predominantly contain the sequences from positive datasets, i.e. the molecules that have high affinity binding to the target. We hypothesize that such clusters are more likely to contain functionally important sequences, which could be of interest for more detailed further investigations in experiments. To identify which sequences belong to clusters and to extract those sequences, we added the clustering analysis functionality into our Jupyter notebook codes for analyses of datasets of affinity-selected molecules.

**Figure 4** shows example clustering analyses using the t-SNE map of sequences containing C*CATTCCGCT motif from SDNA-pos and SDNA-neg datasets. The same map is shown with the labeling of clusters obtained by four different methods, including DBSCAN, Birch, Gaussian Mixture, and k-means methods. When performing the clustering analyses, all four methods required tuning of hyperparameters to improve the labeling of clusters. Among four tested method, the best labeling of clusters was obtained with the DBSCAN method (**Figure 4a**), using ε = 6 and 20 as the minimum number of points forming a cluster. Birch and Gaussian Mixture methods performed well when labeling small clusters, but poorly when labeling the big cluster. Our tuning the hyperparameters for Birch and Gaussian mixture methods did not improve the clustering results. The labeling of clusters by the k-means method was also worse than by the DBSCAN method, since one visually distinct large cluster is labeled as multiple clusters, and adjustments in the number of clusters lead to mislabeling of small clusters, as seen in the bottom left part of the plot in **Figure 4d**. After the clustering analysis, the code has an option to output the sequences that form a specific cluster of interest in the csv format file.

**Figure 4.**
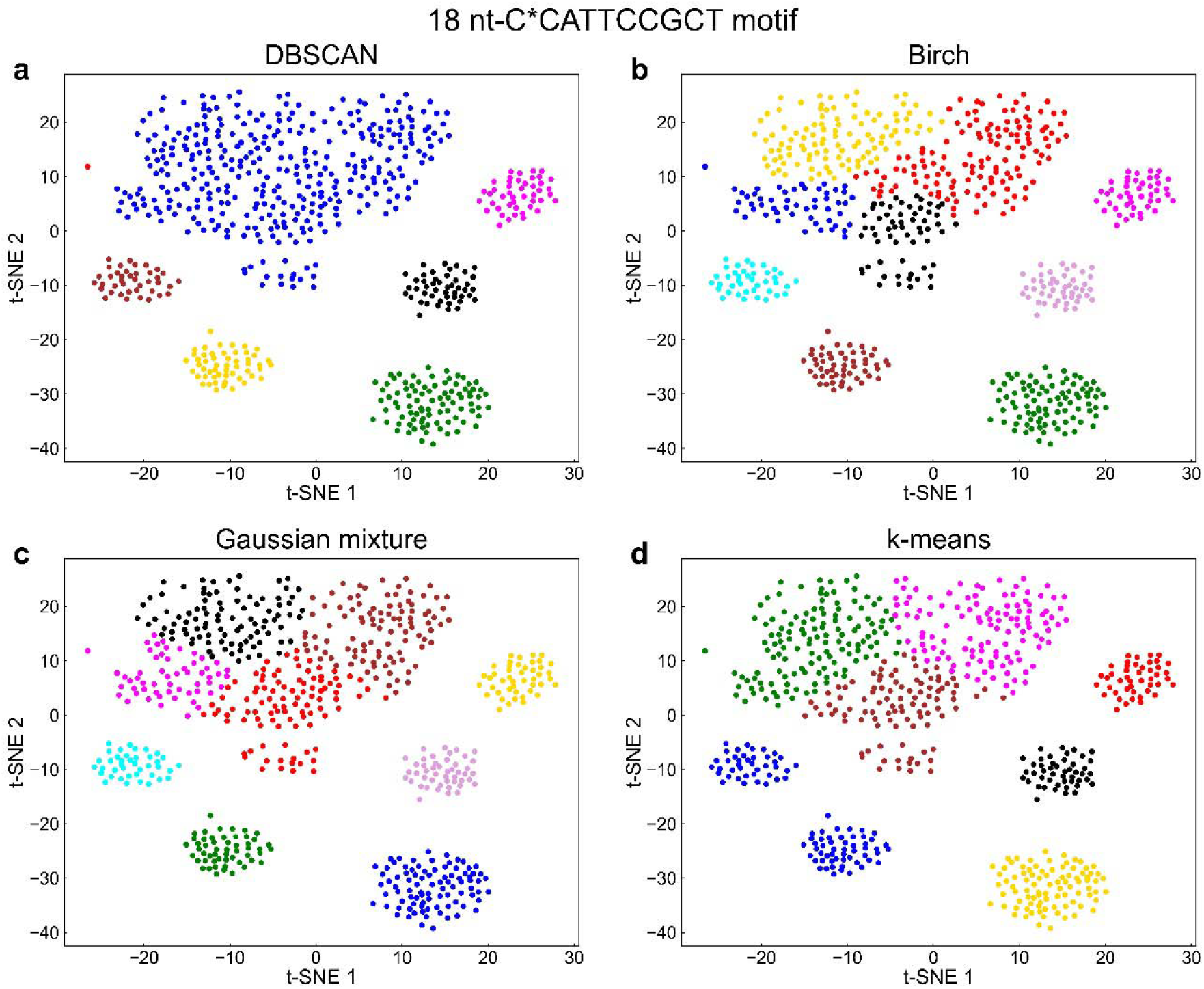
Clustering analysis of the t-SNE map of sequences containing the C*CATTCCGCT motif in the SDNA dataset. The clustering analysis is performed on the two-dimensional t-SNE map described and shown in Figure 3c. a) Clustering analysis using the DBSCAN method, with ε = 6 and 20 as the minimum number of points forming a cluster. b) Clustering analysis using the Gaussian Mixture method, with settings of 9 clusters, spherical covariance type, and 5000 iterations. c) Clustering analysis using the BIRCH method, with settings of 9 clusters. d) Clustering analysis using the k-means method, with settings of 7 clusters.

## Conclusion

The BinderSpace package is a tool for analyzing datasets of sequences of DNA and peptide-based molecules, obtained in selection experiments, such as SELEX^13,14^ and phage display^18^, among others. Our open-source package, implemented in Python3, can perform motif analysis, sequence space visualization, cluster analyses, and sequence extraction from clusters of interest. One of the key features of BinderSpace is its motif search task, based on the Apriori algorithm, which works even for datasets of very short sequences. The motif analysis, providing text-based and visual output, can also be accompanied by visual maps of user-defined functional properties for all the motif-containing molecules for which the functional property was previously measured. Users can also run PCA and t-SNE analyses on their datasets, which can be performed on whole datasets and on motif-related subsets of the data. Functionally important sequences can also be highlighted in the resulting PCA and t-SNE maps. If points (sequences) in two-dimensional maps in PCA or t-SNE space form clusters, users can perform the clustering analyses on their data. Overall, BinderSpace can be used to investigate the relationships between molecule sequences and mechanisms of molecular recognition of the binding targets by these sequences.

## Acknowledgments

We gratefully acknowledge Sanghwa Jeong, Markita P. Landry, Arunika Ekanayake and Ratmir Derda for sharing datasets for testing the performance of our codes. We acknowledge the support of the NSF CBET-2106587 award and the computer time provided by the Texas Advanced Computing Center (TACC).

