## Supplementary Information for "*BinderSpace*: A Package for Sequence Space Analyses for Datasets of Affinity-Selected Oligonucleotides and Peptide-Based Molecules"

Electronic Supplementary Information (ESI)

**Table S1.** Example of encoding method.
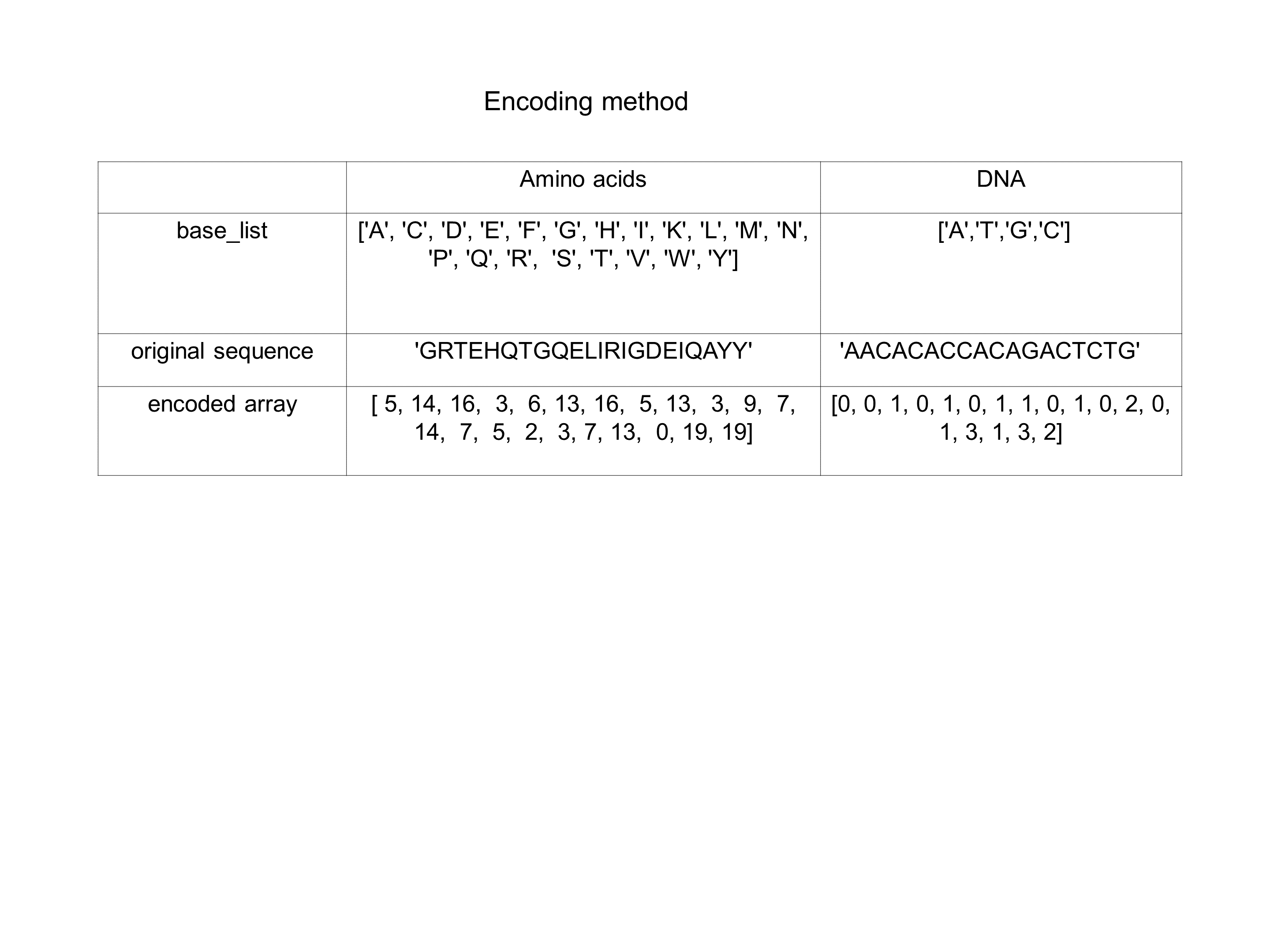

**Table S2.** Example running times for motif search in BinderSpace (binderspace_motif.py) for different datasets and motif search settings.

| Dataset | Frequency | Number of gaps | Running time |
| --- | --- | --- | --- |
| SDNA  (18-nt DNA sequences, positive: 570925, negative : 219381) | 0.001 | 1 | 1128s |
|  | 0.01 | 1 | 85s |
|  | 0.01 | 0 | 25s |
| BCA | 0.1 | 2 | 2.03 |
|  | 0.1 | 0 | 0.74 |

**Table S3.** Examples of motifs found using BinderSpace for SDNA example dataset.

| **SDNA motifs (11-nt with gap)** | **occurrences in positive set** | **occurrences in positive set** | **percentage in positive set** | **percentage in negative set** | **difference** |
| --- | --- | --- | --- | --- | --- |
| CACCGATCCT* | 792 | 34 | 0.1387 | 0.0155 | 0.1232 |
| ACACCGATCC | 828 | 72 | 0.1450 | 0.0328 | 0.1122 |
| ACCGATCCT*C | 708 | 34 | 0.1240 | 0.0155 | 0.1085 |
| ACCGATCCTA* | 645 | 28 | 0.1130 | 0.0128 | 0.1002 |
| ACCGATCCTC* | 628 | 24 | 0.1100 | 0.0109 | 0.0991 |
| **SDNA motifs (10-nt with gap)** | **occurrences in positive set** | **occurrences in positive set** | **percentage in positive set** | **percentage in negative set** | **difference** |
| ACCGATCCT* | 2021 | 100 | 0.3540 | 0.0456 | 0.3084 |
| CACCGATCC* | 2031 | 172 | 0.3557 | 0.0784 | 0.2773 |
| ACCGATCC*A | 1721 | 116 | 0.3014 | 0.0529 | 0.2486 |
| C*CCGATCCT | 1534 | 91 | 0.2687 | 0.0415 | 0.2272 |
| A*CCGATCCT | 1507 | 86 | 0.2640 | 0.0392 | 0.2248 |
| **SDNA motifs (9-nt with gap)** | **occurrences in positive set** | **occurrences in positive set** | **percentage in positive set** | **percentage in negative set** | **difference** |
| ACCGATCC* | 5193 | 446 | 0.9096 | 0.2033 | 0.7063 |
| CCGATCCT* | 4217 | 259 | 0.7386 | 0.1181 | 0.6206 |
| CCGATCC*A | 3818 | 284 | 0.6687 | 0.1295 | 0.5393 |
| A*CCGATCC | 3782 | 297 | 0.6624 | 0.1354 | 0.5270 |
| CCGATCCA* | 3783 | 333 | 0.6626 | 0.1518 | 0.5108 |
| **SDNA motifs (9-nt without gap)** | **occurrences in positive set** | **occurrences in positive set** | **percentage in positive set** | **percentage in negative set** | **difference** |
| ACCGATCCT | 2295 | 119 | 0.4020 | 0.0542 | 0.3478 |
| CACCGATCC | 2033 | 175 | 0.3561 | 0.0798 | 0.2763 |
| CCGATCCTA | 1488 | 63 | 0.2606 | 0.0287 | 0.2319 |
| CATTCCGCT | 1637 | 124 | 0.2867 | 0.0565 | 0.2302 |
| ACCGATCCA | 1512 | 112 | 0.2648 | 0.0511 | 0.2137 |
| **SDNA motifs (8-nt with gap)** | **occurrences in positive set** | **occurrences in positive set** | **percentage in positive set** | **percentage in negative set** | **difference** |
| CCGA*CCT | 7039 | 647 | 1.2329 | 0.2949 | 0.9380 |
| CC*ATCCT | 6908 | 645 | 1.2100 | 0.2940 | 0.9160 |
| ACC*ATCC | 7398 | 901 | 1.2958 | 0.4107 | 0.8851 |
| CCG*TCCT | 6469 | 614 | 1.1331 | 0.2799 | 0.8532 |
| ACCGA*CC | 7143 | 906 | 1.2511 | 0.4130 | 0.8382 |
| **SDNA motifs (8-nt without gap)** | **occurrences in positive set** | **occurrences in positive set** | **percentage in positive set** | **percentage in negative set** | **difference** |
| CCGATCCT | 4881 | 301 | 0.8549 | 0.1372 | 0.7177 |
| ACCGATCC | 5198 | 449 | 0.9105 | 0.2047 | 0.7058 |
| CCGATCCA | 3982 | 347 | 0.6975 | 0.1582 | 0.5393 |
| TCCGATCC | 2910 | 254 | 0.5097 | 0.1158 | 0.3939 |
| CGATCCTA | 2455 | 167 | 0.4300 | 0.0761 | 0.3539 |

**Table S4.** Examples of motifs found using BinderSpace for BCA example dataset.

| **BCA motifs (4-aa with 2 gaps)** | **occurrences in positive set** | **occurrences in positive set** | **percentage in positive set** | **percentage in negative set** | **difference** |
| --- | --- | --- | --- | --- | --- |
| HW** | 47 | 21 | 0.6014 | 0.2747 | 0.3267 |
| H**V | 48 | 24 | 0.6142 | 0.3140 | 0.3002 |
| H*Y* | 45 | 22 | 0.5758 | 0.2878 | 0.2880 |
| H**Y | 45 | 22 | 0.5758 | 0.2878 | 0.2880 |
| HV** | 45 | 22 | 0.5758 | 0.2878 | 0.2880 |
| **BCA motifs (3-aa with 1 gap)** | **occurrences in positive set** | **occurrences in positive set** | **percentage in positive set** | **percentage in negative set** | **difference** |
| HV* | 81 | 47 | 1.0365 | 0.6149 | 0.4216 |
| V*W | 81 | 47 | 1.0365 | 0.6149 | 0.4216 |
| H*W | 82 | 48 | 1.0493 | 0.6279 | 0.4213 |
| WD* | 91 | 61 | 1.1644 | 0.7980 | 0.3664 |
| MH* | 62 | 34 | 0.7933 | 0.4448 | 0.3486 |
| **BCA motifs (3-aa without gap)** | **occurrences in positive set** | **occurrences in positive set** | **percentage in positive set** | **percentage in negative set** | **difference** |
| TLT | 9 | 0 | 0.1152 | 0 | 0.1152 |
| EVV | 8 | 0 | 0.1024 | 0 | 0.1024 |
| SSY | 8 | 0 | 0.1024 | 0 | 0.1024 |
| KWD | 9 | 1 | 0.1152 | 0.0131 | 0.1021 |
| FVH | 10 | 2 | 0.1280 | 0.0262 | 0.1018 |

| **BCA motifs (2-aa without gap)** | **occurrences in positive set** | **occurrences in positive set** | **percentage in positive set** | **percentage in negative set** | **difference** |
| --- | --- | --- | --- | --- | --- |
| VH | 148 | 105 | 1.8938 | 1.3736 | 0.5202 |
| KV | 115 | 74 | 1.4715 | 0.9681 | 0.5034 |
| WD | 133 | 93 | 1.7019 | 1.2166 | 0.4852 |
| VV | 103 | 66 | 1.3180 | 0.8634 | 0.4546 |
| HY | 110 | 73 | 1.4076 | 0.9550 | 0.4526 |

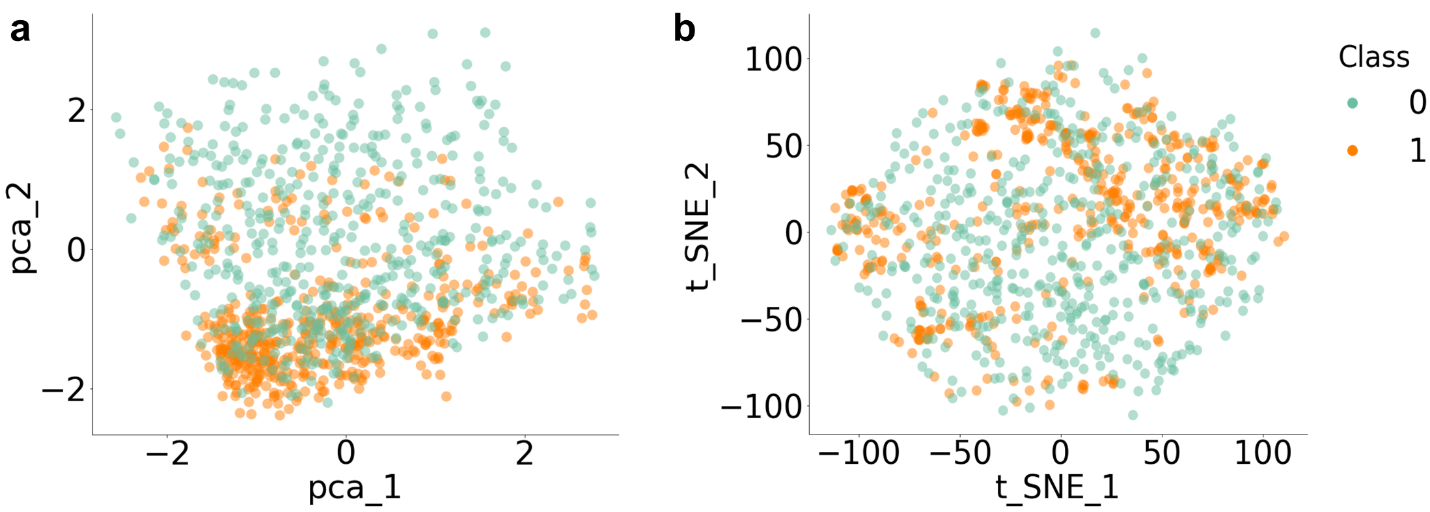

**Figure S1.** Example output of simple PCA and t-SNE analyses, which can be plotted it in 2- and 3-dimensional figures with BinderSpace. The example is showed for analyses of complete SDNA datasets, where only the top 500 sequences from SDNA-pos and top 500 sequences from SDNA-neg are shown.
